# Determining epitope specificity of T-cell receptors with transformers

**DOI:** 10.1101/2023.03.31.534974

**Authors:** Abdul Rehman Khan, Marcel JT Reinders, Indu Khatri

## Abstract

**Motivation:** T-cell receptors (TCR) on T cells recognize and bind to epitopes presented by the major histocompatibility complex (MHC) in case of an infection or cancer. However, the high diversity of TCRs, as well as their unique and complex binding mechanisms underlying epitope recognition, make it difficult to predict the binding between TCR and epitope. Here, we present the utility of transformers, a deep learning strategy that incorporates an attention mechanism that learns the informative features, and show that these models pretrained on a large set of protein sequences outperform current strategies.

**Method:** We compared three pre-trained auto-encoder transformer models (ProtBERT, ProtAlbert, ProtElectra) and one pre-trained auto-regressive transformer model (ProtXLNet) to predict the binding specificity of TCRs to 25 epitopes from the VDJdb database (human and murine). Two additional modifications were performed to incorporate gene usage of the TCRs in the four transformer models.

**Results:** Of all 12 transformer implementations (4 models with 3 different modifications), a modified version of the ProtXLNet model could predict TCR-epitope pairs with the highest accuracy (weighted F1 score 0.55 simultaneously considering all 25 epitopes). The modification included additional features representing the gene names for the TCRs. We also showed that the basic implementation of transformers outperformed the previously available methods, i.e. TCRGP, TCRDist and DeepTCR, developed for the same biological problem, especially for the hard-to-classify labels.

**Conclusion:** We show that the proficiency of transformers in attention learning can indeed be made operational in a complex biological setting like TCR binding prediction. Further ingenuity in utilizing the full potential of transformers, either through attention head visualization or introducing additional features, can further extend T-cell research avenues.

**Availability:** Data and code are available on https://github.com/InduKhatri/tcrformer

## 1. Introduction

The human immune system can mount the immune response by generating multiple T-cell receptors (TCRs) in response to a pathogenic infection. Principally, this response involves interaction between T-cell receptors (antigen/epitope-recognition receptors on T cells) and the epitopes (short peptides from pathogenic proteins) present in infectious agents (bacteria/viruses). Complementarity determining region 3 (CDR3) on both α and β chains of TCRs bind with the epitope. The diversity of TCRs is estimated to be approximately 10^18^ in humans and 10^15^ in mice. The hypervariability of the CDR3 region is imparted during the V(D)J recombination process followed by the junctional diversity due to the addition of additional bases during the recombination process (see Supplementary text for more detail). Determining the binding specificity of such highly diverse and variable TCR sequences to an epitope is a challenging problem since multiple TCRs can bind to an epitope and similar antigenic peptides can be recognized by a multitude of TCRs. Learning the specificity of TCRs to the epitopes will enhance our understanding of the generation of the immune responses at the receptor level to multiple similar epitopes and/or different epitopes from different infectious agents.

An alignment model or *attention* focuses a neural network on learning relevant relationships, reducing heavy translations and improving translation performance. One of the deep neural architectures using attention is transformers [1]. Unlike long short-term memory (LSTM) models, transformers are not restricted by the input sequence length. transformers are trained using Transfer Learning, in which transformer models are first pre-trained on task-analogous objectives and fine-tuned on task-oriented objectives (see Supplementary text for more detail). Through transfer learning, a transformer can learn contextual information from a large dataset during pre-training and then apply the knowledge learnt from pre-training towards a downstream task, which generally has a scarcity of data. Since the introduction of transformers, many variations of its architecture have been released [2–5]. Common in all these transformer architectures is the *self-attention* mechanism. While attention focuses on parts of the input that leads to a better conclusion, self-attention will focus on surrounding words to delineate the meaning between similar words. Furthermore, multi-headed self-attention learns multiple representations, each of these are termed as a head [6]. Different transformer models implement different strategies to utilise attention mechanisms more efficiently.

Remarkably, transformers are not just limited to natural languages. Given a sufficient large corpus, they can be tailored towards any sequence-based inference. ProtTrans is a transformer model trained on a large corpus of protein data, e.g. the Uniref and BFD databases, comprising 216 million and 2.122 billion protein sequences, respectively, and has been shown to learn the contextual grammar of amino-acid sequences [4]. Analogous to NLP tasks, the ProtTrans pre-training task involved a vocabulary of twenty amino acids. Therefore, sentences and words are analogous to protein sequences and amino acids, respectively. Four of variants of this ProtTrans transformer model, three auto-encoders (ProtBERT [6], ProtAlbert [5], ProtElectra [2]) and an auto-regressive model (ProtXLNet [3]), are publicly available as of November 2021. It has been shown that their attention mechanism to find relevant regions in a protein sequence (pattern of amino acids) accurately emulates knowledge needed to understand the mechanisms behind a protein function. This inspired us to use transformers to learn the patterns of a TCR-epitope interaction to further our understanding of the processes determining the affinity between a TCR and an epitope.

This work demonstrates the feasibility of transformers to predict the TCR specificity to individual epitopes based on TCR sequence information. We show how to adapt the pre-trained transformers to the setting of predicting TCR-epitope binding specificity. We compare performances for the different transformer models (ProtBERT, ProtAlbert, ProtElectra and ProtXLNet), and we compare their performance to previously available tools based on distance metrics [7] or deep learning [8]. It is known that additional information about a TCR (besides their sequence), such as gene name, MHC class can enhance the predictive power of the models [7,8]. However, transformers are known to accept only sequence data as input. We uniquely modified the publicly available transformers to incorporate information about TCR gene names, as this information was shown to impart specificity to recognize a epitope [9]. All together, we show that the use of transformers to predict TCR binding specificity to unique epitope is very promising.

## 2. Methods

### 2.1 Data acquisition and preparation

VDJdb provides a centralised source for TCR-epitope pairs. For a given TCR sample, the database provides CDR3, V and J sequences of the TCRβ and/or TCRα receptor protein, the MHC Class I/II annotation, as well as the organism in which it was observed [10]. For epitopes, the database provides the epitope sequence, the parent gene of the epitope, and the antigen species for the epitope. Confidence scores (0, 1, 2 and 3) are associated with each TCR-epitope pair, wherein a zero score indicates computationally predicted specificities, and higher scores (i.e. 1, 2, and 3) represent pairs that were validated using one or more wet-lab based techniques, e.g., assay identification, TCR sequencing, or verification procedure.

Data obtained from VDJdb consisted of 81,762 entries (as per November 2021), which included epitopes binding to a variety of antigens from cancer, immune-disorders, plants, microbes etc. We filtered the dataset by selecting TCR-β chains of Human and Murine epitopes with a confidence score greater than 0. Only single instances of duplicate CDR3 sequences with the same V-gene, J-gene, MHC A, and MHC B were retained. Additionally, we removed epitope species not originating from infectious agents, i.e. related to autoimmune disorders, cancer, synthetic, bacterial antigens or allergies. Finally, we only kept the TCR-epitope pairs (called “classes” from here of) comprising of minimum 50 CDR3 sequences belonging to the TCR-epitope pair. **Table 1** summarises the number of TCR-epitope pairs left after the different filter steps. Finally, 2.674 pairs have been used for training, validating and testing purposes. This consisted of 25 classes unique TCR-epitope pairs (classes): , 10 classes with more than 100 instances (these were considered “Easy-to-classify”) and 15 classes with less than 100 samples (these were considered “Hard-to-classify”) (**Supplementary Figure 1**). Information about unique V and J genes (63 V genes and 13 J genes) was encoded using an ordinal encoder and associated with each TCR sequence. Consequently, input data for each TCR-epitope pair consists of the collection of the Epitope’s species, genes, and sequence.

**Table 1:** Number of TCR-epitope pairs retrieved from the VDJdb database after each filtering step. *Duplicates were removed based on the same gene and MHC gene values and CDR3 sequence

| Filtration criteria | Sample count |
| --- | --- |
| Raw data | 81762 |
| Post removing N/A | 80679 |
| Selecting only human and murine antigen | 78647 |
| Selecting TCR-epitope pairs with confidence score > 0 | 9580 |
| Post removing allergen and cancer antigen | 8547 |
| Post removing duplicated entries* | 5680 |
| Selecting the TCR-epitope pairs with TRB sequences | 4057 |
| Selecting the TCR-epitope pairs with >=50 samples | 2674 |

### 2.2 Multi-class classification of TCR-epitope pairs

Four different models i.e. ProtBERT [6], ProtAlbert [5], ProtElectra [2] and ProtXLNet [3] were used for learning and predicting the TCR-epitope pairs in a multi-class setting [4]. We used pre-trained models. The ProtBERT, ProtAlbert, and ProtXLNet models were pre-trained on the Uniref database [11], and ProtElectra was trained on the Big Fantastic Database (BFD) database (https://bfd.mmseqs.com/). These pre-trained models were subsequently used to learn to predict TCR-epitope specificity. Hereto the 2.674 TCRs divided over 25 different classes of epitopes is split into 70% training, 15% validation, and 15% testing dataset in a stratified way.

#### Using pre-trained transformers to predict TCR-epitope specificity

Each transformer has its specific tokenizer, which prepares the sequence-based input using its pre-trained vocabulary. A general overview of the transformer model is depicted in **Figure 1A**. The embedding block prepares input for the transformer, which includes adding special tokens to denote special relationships (such as padding), an attention mask to denote if a token is to be considered for attention calculation, and a segment ID to denote two separate sequences in the same input. The tokenizer in the embedding block also performs the encoding of labels and presents sequences to the transformer block. Each tokenizer maintains homogeneity of the sequence encoding and presents each sample to the transformer to learn a representation. The representation of the input sequence is then fed to a classification block, which performs sequence inference tasks. In the baseline transformer (**Figure 1A**), the classification block receives an n-dimensional (for example, in the case of ProtBERT, n=768) representation of each amino acid as input from the final hidden state of the transformer block. The first token of every sequence is a special classification token (signified as CLS by the tokenizer) which contains the aggregated representation of the input sequence in the final hidden state. This representation is termed pooled output. The pooled output is the input to the classification block to perform the TCR-epitope classification task. The standard classification block consists of a linear layer mapping the pooled output vector (*x* of the size of the hidden layer) from the transformer block as shown in Eq. 3 to class label (y) where y is the number corresponding to the classes.

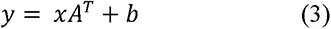

**Figure 1:**
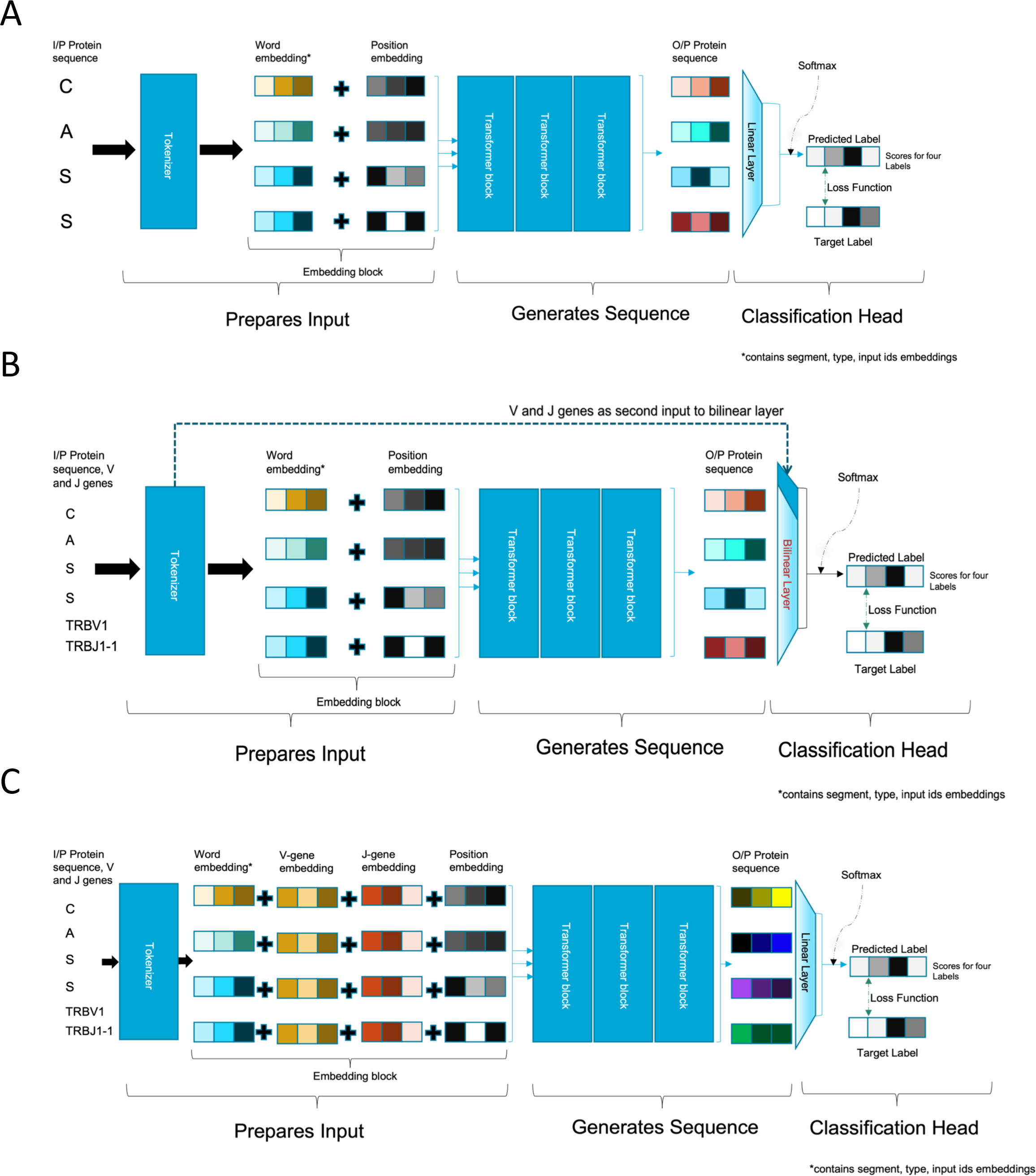
Overview of the data flow in the different Transformers settings. (a) *Baseline setting*. The CDR3 sequence is provided to the tokenizer to generate a numerical representation of the input sequence based on embeddings learnt during pre-training. Prepared input is divided into three parts (Query, Value and Key) and provided to the transformer block to implement the self-attention mechanism. The output representation generated by the transformer block is then used by the classification block to predict the epitope label. (b) *Classification setting*. Sequence is provided to the transformer block and the gene usage information is directly presented to classification head, where the gene information is added into a bilinear layer. (c) *Embedding setting*. The sequence along with the gene usage information from the embedding block is provided to transformer block, thereafter the combined embeddings are is fine-tuned.

#### Loss Function for Imbalance data

The imbalance in our dataset prompted us to adapt the loss function. A binary loss function created upon cross-entropy is Focal-loss, [12] which adds a modulating factor (or Focusing parameter) to cross-entropy. The modulating factor is adjusted using gamma (γ > 0), which reduces the contribution of samples from the “easy-to-classify” classes, and enhances the contribution of samples of the “hard-to-classify” classes to the loss value. The enhanced loss value of samples from the hard-to-classify çlasses informs the model to accommodate more for these classes, which otherwise is overlooked during training. The loss appears normal to the model; the adjusted loss value is the manipulation performed before providing it to the model, and the model can then direct the gradient accordingly.

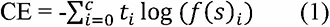

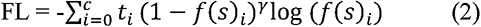

Multi-class cross-entropy can be extended to multi-class focal loss with an additional step after calculating the sum of losses for each class, i.e., applying the modulating factor as shown in Eq. 2. A recommendation of γ = 2 was made by the authors of focal loss; however, they only assumed binary cases. For the multiclass classification model in our problem, we used γ as hyperparameter to optimise.

#### Adding non-sequence information to the classification task

We sought ways to include information about gene names as additional input as this has shown to improve TCR-epitope binding specificity prediction performance. To avoid re-training the transformer models (which would take an enormous amount of time, see Supplementary information: Transfer Learning), we altered the tokenizer to accommodate additional features along with sequences. We explored two different ways to do that: 1) before, and 2) after the transformer blocks. The first option requires a modification in the classification block of the model (**Figure 1B**). The second option requires a modification in the embedding block (**Figure 1C**).

#### Adding additional features by modifying the classification block

The classification block receives the additional features directly from the tokenizer, bypassing the transformer block (**Figure 1B**). Each CDR3 sequence has V and J gene names associated with it. An ordinal encoding for these gene names is provided to the tokenizer, which it associates with each sample and then given to the classification block. We replaced the linear layer with a bi-linear layer where the pooled output (*x*_1_ having a size equal to the size of the hidden layer) and gene encoding (*x*_2_ with size 2 representing V and J gene names separately) is mapped to a class label (*y*). The bi-linear transformation, as shown in Eq. 4, will calculate weight matrix (*A*) and bias (*b*) based on the two input vectors (sequence, *x1*, and gene name information, *x2*), which will express the interaction between sequence and gene names.

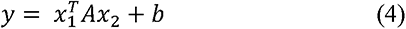

#### Adding additional features by modifying the embedding block

The embedding block of a pre-trained model contains a learnt representation of its vocabulary. The introduction of additional information would require learning new representations. To avoid this, we propose to add two embedding layers; one for all unique V genes, and another for all unique J genes (**Figure 1C**). The padding index for these layers is different from the padding for the word embedding layer (so we explicitly pass an alternative padding index for the additional features; 0 indices for V and J genes embedding layer). Special tokens were added before and after a protein sequence during tokenization. The separate padding index will associate a gene embedding to only a protein sequence and not to the special tokens. Each sequence embedding and the V and J gene names are then merged. The new representations were presented to the classification block.

The padding index, the unique genes names and the special token together, resulted in a total embedding size for V-genes of 65, and for J genes of 15. These new layers were randomly initialised when fine-tuning. To synchronize with the fine-tuning of the rest of the model, these weights are learned at an increased rate (for the sake of simplicity, by a factor of 10).

### 2.3 Hyperparameter optimisations and performance evaluation

A Bayesian algorithm (Tree Parzen Estimator) is utilised to optimise the hyperparameters of the different models using Optuna [13], which is recommended for situations that require the exploration of many hyperparameters. Experiments are tracked using comet.ml and PyTorch for training each model. Optimization is done for ten major hyperparameters: gradient accumulation, learning rate, weight decay, attention layer dropout (not included in auto-regressive or AR model), hidden layer dropout (‘dropout’ in AR model), classifier layer dropout (‘summary last dropout’ in AR model), adam_beta1, adam_beta2, warmup ratio, and gamma. A seed as a hyperparameter ensures model stability, but performance is not evaluated on seed values.

When optimizing hyperparameters we gain insight into the behaviour of the transformers when trained with TCR data. Optuna [13] provides sampling strategies depending on the hyper-parameter of choice. Table 2 presents the sampling strategy for each hyper-parameter and its range. Since training steps are directly dependent on training batch sizes, and since we have small training data of 1871 samples, we used a training batch size and an evaluation batch size of 1 and 8, respectively. In preliminary experiments, we experienced high fluctuations in evaluation loss, which made transformer training susceptible to stopping prematurely. Therefore, each experiment is trained with an early stopping for 50 training epochs giving sufficient training time to adjust fluctuations caused by focal loss.

**Table 2:** The list of tested hyperparameters with range of considered values (“modulations”).

| Hyperparameters | Modulations |
| --- | --- |
| Gamma | (0, 10) with step of 0.5 |
| Adam beta 1 | [0.5, 0.9] with step of 0.01 |
| Adam beta 2 | [0.5, 0.99] step of 0.001 |
| Learning rate | [10<sup>-5</sup>, 10<sup>-2</sup>] from log domain |
| Weight decay | 0, 0.01, 0.001, 0.0001, 0.00001, 0.000001 |
| Gradient accumulation steps | [1, 128] |
| Classifier / Summary layer dropout | [0.5, 0.9] with step of 0.1 |
| Hidden layer / dropout | [0.5, 0.9] with step of 0.1 |
| Attention layer dropout | [0.5, 0.9] with step of 0.1 |
| Warmup ratio | 0.10, 0.20 |
| Seed (for model stability) | [1, 100] |

#### 2.3.1 Evaluation metrics

We used the weighted F1-score and ROC plots as metrics to assess the predictability of the transformer models. Additional metrics included were a balanced accuracy score, weighted precision, and weighted recall to provide an overview of the different performances of a model. Parallel coordinate plots and parameter importance were plotted to evaluate the contribution of parameters towards the objective value.

In a multi-class setting, the assessment of transformers is done through the epitope-specific perspective and model-specific perspective. The area under the ROC curve provides an epitope-specific perspective where performance comparison for TCR specificity can be made. The weighted F1-score of the model provides us with a holistic measure for comparing different models (across all methods) in a multi-class setting and hence a model-specific perspective.

The performances of all 12 transformer implementations (4 transformer models each with 3 different modifications) were evaluated using ROC plots. The area under ROC (AUROC) curve for all 25 labels were assessed using the Wilcoxon test to compare performance across different implementations of the same transformer model. The transformer implementation with no modifications for each model are called baseline while the ones with modifications in classification and embedding blocks are called classification and embedding.

We compared the transformer models to existing (non-transformer) models. TCRGP and TCRDist both have 24 common epitopes that overlap with our filtered dataset. DeepTCR has 20 epitopes in common. We used the AUC values published in these studies to compare with the AUC values of our models.

## 3. Results

We compared the performance of four different transformer models, three of which are based on an autoencoder architecture (ProtBERT, ProtAlbert, ProtElectra) and one is based on an auto-regressive structure (ProtXLNet), as well as their modification to include information about the V/J genes, on a dataset containing binding specificities between 2.674 TCR-epitope pairs. These pairs are split in 25 classes, with 10 classes having more than 100 instances (and considered “Easy-to-classify”), while 15 classes have less than 100 samples (and considered “Hard-to-classify”) (**Supplementary Table 1**). Data was split in 70% training, 15% validation, and 15% testing dataset in a stratified way across the classes. The validation set was used to optimize the hyperparameters, and the testing set was used to estimate the performance of the optimized models. Each of the four different transformer models were run in three different settings: 1) without the use of information about the V/J genes (*Baseline setting*), 2) including the V/J gene names within the classification block (*Classification setting*), and 3) including information about the V/J names by modifying the embedding layer (*Embedding setting*). See Figure 1 and Methods for more details. This resulted in 12 optimized transformer models (four different transformers, in three different settings). These optimized models were subsequently compared to three of the current best-performing tools classifying TCR-epitope pairs, i.e. TCRGP, TCRdist and DeepTCR (all non-transformer-based). We first discuss the tuning of the hyperparameters and the learning behaviour of the transformer models. Then we compare the classification performances between the different models.

### 3.1 Learning behaviours of the transformer models

The three auto-encoder models had eleven hyperparameters (including the seed value), while the auto-regressive model had ten hyperparameters (**Table 2**). Optimization for these hyperparameters was performed for at least 100 runs for each implementation. For the ProtBERT and ProtElectra autoencoder models the hidden-layer dropout probability had the largest influence on the optimization (65-87% and 71-81%, respectively) (**Figure 2**). Moreover, for both models the importance of the hyperparameters is roughly the same across the three different settings. For the ProtAlbert autoencoder, we observe a more equal influence across hyperparameters. Within the baseline setting, the hidden-layer dropout probability is still most influential although the learning rate is also important, while in the other settings the learning rate is the most influential (21-39%). This could be because ProtBert and ProtElectra do not have parameter sharing among the encoder blocks, resulting in a large network size (thus more parameters to train, emphasizing the need for regularisation), whereas ProtAlbert does have parameter sharing among the encoder blocks, resulting in a smaller network size and thus less need for regularisation. As ProtAlbert is not solely dependent on the encoding representation, but also on the learning rate, it might be more efficient in learning new relationships from the interaction between TCRs and epitopes.

**Figure 2:**
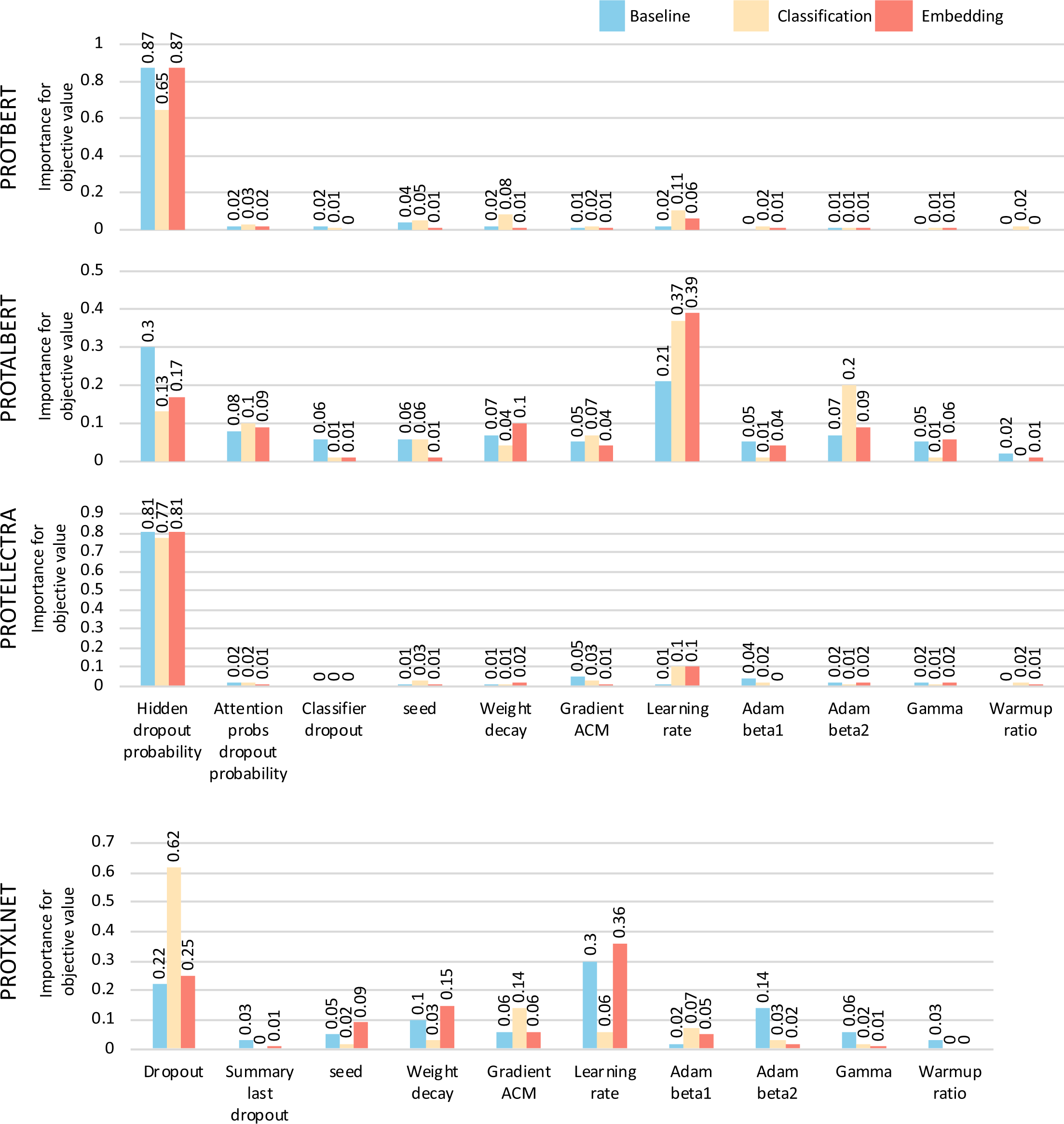
Performance of the hyperparameters for all considerd transformer models. The bar graphs are colored based on the methods implemented on the different transformer models.

The overall importance of the hyperparameters in the Classification setting of the ProtXLNet autoregressive model was similar to the ProtBert and ProtElectra auto-encoders, with the dropout probability being most influential (62%). The Baseline and Embedding settings for ProtXLNet showed a behaviour which was more similar to ProtElectra, with the learning rate being most important (30% and 36%, respectively). From this we conclude that ProtXLNet finds it more difficult to introduce the gene usage in the classification block, as it is trying to generate a better representation from the transformer block.

### 3.2 Comparing performances of transformer models

When comparing the weighted F1 scores of the transformer models, we see a general trend in which ProtXLNet performs better than ProtElectra, which in turn performs better than ProtAlbert. ProtBert performs the worst (**Figure 3A, Supplementary Table 2**). Moreover, we see a general trend that the Embedding setting performs best, followed by the Baseline setting. The Classification setting performed worst, although for ProtBert, the classification setting performs best. On contrary, we observed that the Embedding setting outperformed the other settings for the rest of the transformer models, i.e. ProtAlbert, ProtElectra and ProtXLNet. In this setting the number of unclassified and misclassified labels reduced significantly (**Supplementary Figure 2**).

**Figure 3:**
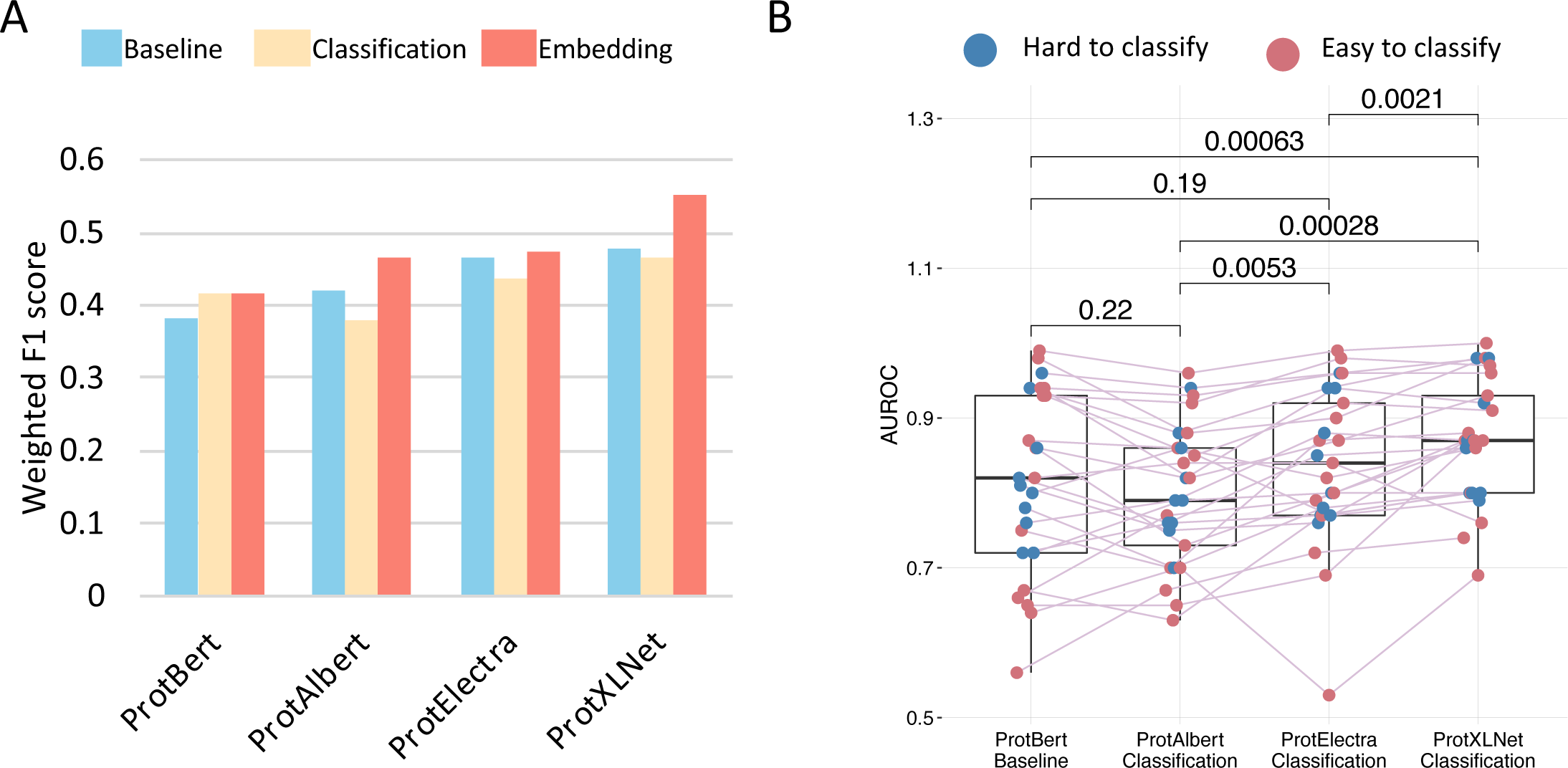
Comparison metrics for all transformer models. (a) weighted F1-score on test dataset, (b) AUCROC scores for the different classes in the test dataset. Only the significant P values as calculated by Wilcoxon paired test are mentioned in the plot.

When comparing the mean AUCROC scores of all classes within the TCR-epitope dataset across the best (F1 score) setting for each of the four transformer models, we observe that the AUCROC of the Classification setting of ProtBERT improves performance over ProtAlbert in the Embedding setting, which then becomes the worst. The Embedding setting of the ProtXLNet consistently performed significantly better than the other methods (Wilcoxon paired comparison, all three p-value’s < 0.0021), and has the least unclassified and misclassified labels (**Supplementary Figure 3**).

The 25 classes in the dataset could be divided into ““easy-to-classify” classes and “hard-to-classify” classes (Methods). Indeed, we see that some that some epitopes are hard to classify by all models, for example the CMV and EBV epitopes (**Supplementary Figure 3**). Some models have problems to recognize specific classes, for example the Baseline setting of ProtBert cannot identify seven classes (**Supplementary Figure 3**), indeed belonging to the “hard-to-classify” category, indicating a too low number of training samples for these classes. When investigating the AUCROC scores for the different categories of classes (**Figure 3B**), we see that there is hardly any difference between the AUCROC scores for the ““easy-to-classify” classes and “hard-to-classify” classes for ProtAlbert, ProtElectra and ProtXLNet (scores are evenly distributed) (**Supplementary Table 2)**. ProtBERT, however, seems to struggle more with some of the “easy-to-classify” classes. When studying the differences in classification accuracy between the ““easy-to-classify” classes and “hard-to-classify” classes for the different settings for each transformer, we find no significant differences (**Supplementary Figure 4**). Although based on the performance of the models, we observed that the Classification setting is the most economical as it used the least training time when compared with other methods of the models (**Table 3**).

**Table 3:** Performance metrics of the different transformer models across the three different settings (Baseline, Classification, and Embedding setting) on the test dataset.

| PROT transformers | Settings | Duration (hours) | Weighted F1-score | Balanced Accuracy | Mean AUC |
| --- | --- | --- | --- | --- | --- |
| ProtBERT | Baseline | 4.49 | 0.39 | 0.33 | 0.79 |
|  | Classification | 1.99 | 0.41 | 0.37 | 0.8 |
|  | Embedding | 2.21 | 0.41 | 0.35 | 0.79 |
| ProtAlbert | Baseline | 3.16 | 0.42 | 0.37 | 0.79 |
|  | Classification | 0.57 | 0.38 | 0.33 | 0.78 |
|  | Embedding | 3.21 | 0.46 | 0.44 | 0.81 |
| ProtElectra | Baseline | 1.29 | 0.46 | 0.41 | 0.86 |
|  | Classification | 0.9 | 0.44 | 0.41 | 0.8 |
|  | Embedding | 0.99 | 0.47 | 0.4 | 0.84 |
| ProtXLNet | Baseline | 1.57 | 0.48 | 0.44 | 0.81 |
|  | Classification | 1.21 | 0.46 | 0.38 | 0.8 |
|  | Embedding | 1.23 | 0.55 | 0.5 | 0.88 |

### 3.3 Transformers in comparison with publicly available tools

To assess the best-performing transformer model from our study, we selected a few previously available best-performing tools classifying TCR-epitope pairs: TCRGP, TCRdist and DeepTCR. As all the tools used a common database (VDJdb) for assessing the performances, a head-to-head comparison was possible. However, not all the epitopes were present in all the tools, since the database updates the samples in the classes. TCRGP and TCRdist lacked one epitope, whereas with the DeepTCR only 20 epitopes were in common to the epitopes used in our study. These classification tools were compared to our best model; ProtXLNet in the Embedding setting.

Our ProtXLNet model, which had a mean AUCROC of 0.876 across the 24 common epitopes, showed an improvement of almost 5% when compared with TCRGP (AUCROC=0.831) and TCRdist (AUCROC=0.781) (**Figure 4**). For the three “hard-to-classify” labels of the MCMV epitope family, TCRGP and TCRdist had a mean AUCROC of 0.843 and 0.807, respectively, whereas ProtXLNet had a mean AUCROC of 0.946 for the same labels of MCMV epitope family (an improvement of ≥10%) **(Supplementary Table 3)**.

**Figure 4:**
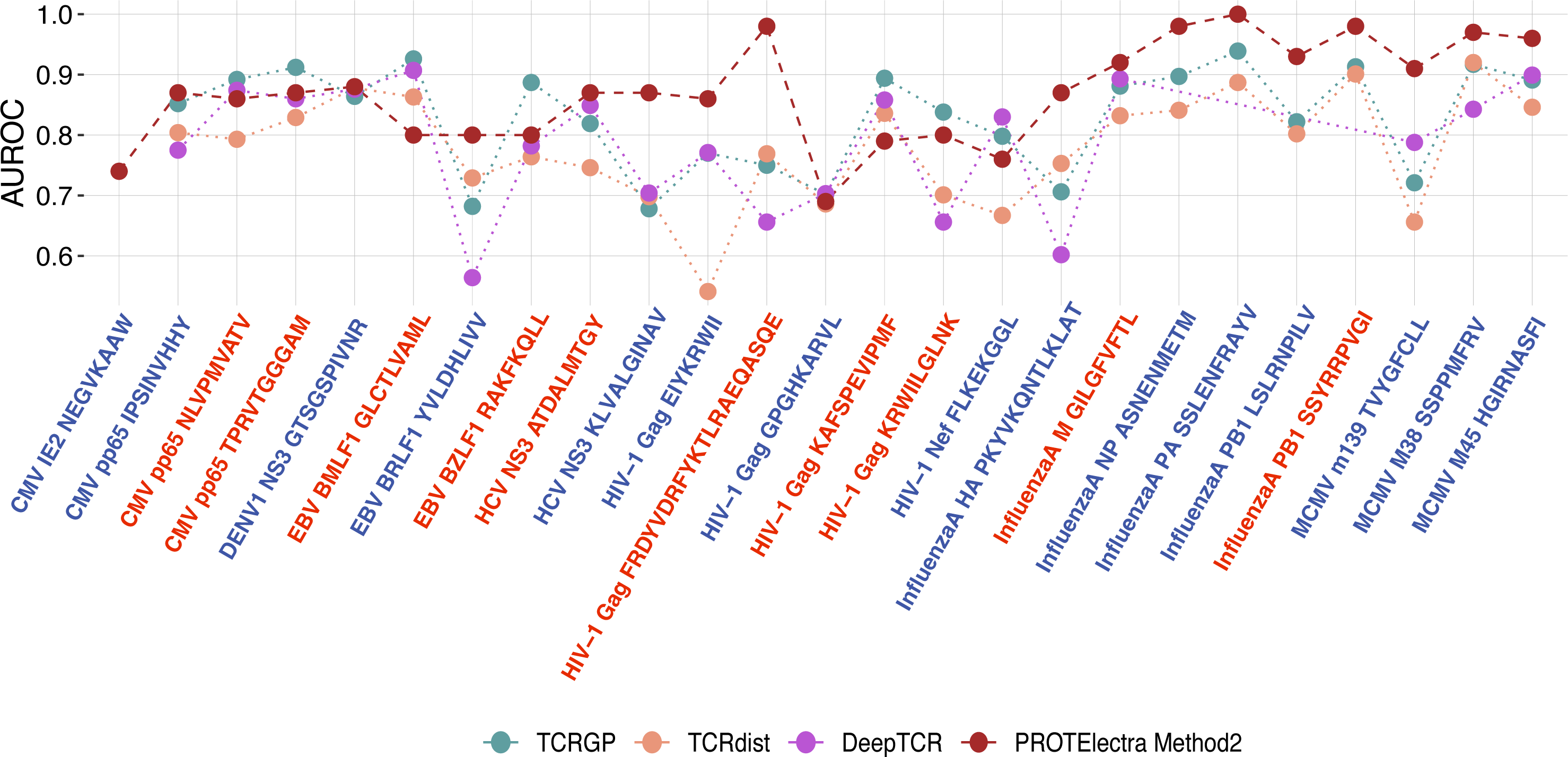
AUCROC scores for the ProtXLNet model with Embedding setting and previously known methods (TCRRGP, TCRdist and DeepTCR) for different epitope classes. The classes are colored blue if the epitope is considered to be of the hard-to-classify class and red for the easy-to-classify class.

Similarly, for the 20 common epitopes with DeepTCR model, ProtXLNet exhibited a mean AUCROC of 0.856, whereas DeepTCR had a mean AUC of 0.784. This difference in the mean AUCROC was mainly attributed to the “hard-to-classify” labels, for which DeepTCR performed poorly (**Figure 4, Supplementary Table 3**).

## 4. Discussion

A diverse set of TCRs are generated by the adaptive immune system that recognizes a short immunogenic peptide on the proteins of the pathogens. The epitopes should ideally have a specific set of TCRs with specific properties, such as specific CDR3 sequences, specific gene usage and a specific MHC class. Several sequence-based and deep learning-based models have been proposed to predict TCR-epitope pairs, however, there is still a performance deficit in these methods. In this study, we used a novel deep learning approach, *transformers*, that uses attention layers that are learnt to focus on relevant parts of the provided TCR sequence. We adopted three pre-trained auto-encoders (ProtBERT, ProtAlbert and ProtElectra) and one auto regressor (ProtXLNet) and added a classification head to build multi-class classification models to identify TCR specificity to the selected epitopes from the VDJdb. To incorporate gene-usage in these transformers, we have made novel modifications to these transformers, either by adapting the classification block (Classification setting), or by adapting the embedding layer (Embedding setting).

Across twelve different transformer models, we find that the ProtXLNet auto-regressor model modified with the Embedding setting performed with the highest weighted F1 score (0.55), the highest Mean AUCROC for the “hard-to-classify samples (0.88) as well as for the “Easy-to-classify” samples (0.86), and significantly better AUC values for all classes (**Supplementary Table 2**). Moreover, this best performing transformer model outperforms previously developed sequence-based (TCRGP and TCRdist) and deep learning (DeepTCR) based methods (**Supplementary Table 3**).

The results can further be evaluated using attention head visualisation in which attention heads playing the most vital role towards a classification can be highlighted. As we can instruct the tokenizer to calculate attention for specific regions in CDR3 sequences, identifying the relevant amino-acid residues from the interaction of TCR and epitope by the attention mechanism can further improve the usage of the transformers for making better predictions of specific TCRs (at the level of amino-acids in the CDR3 sequences) binding to specific epitopes.

The previous methods have shown that incorporating CDR3 sequences and gene usage information of both alpha and beta chains along with MHC restriction can increase the performance and the prediction power of the tools. Here, in our study, we only used the CDR3 sequences and gene usage of the TCR beta chain. We only targeted this simplified approach because our main goal was to understand the different ways to incorporate gene usage within the transformer models (i.e. the Classification setting and the Embedding setting).

During optimization of the different parameters for all four transformer models, we observed that the models tended to stop early for potentially good runs. This might have been caused by the choice of the loss function, as, depending on modulating factor, the focal loss may cause many spikes in loss values. These spiking fluctuations may then be misinterpreted and cause early stopping. Increasing the tolerance threshold for early stopping led to stale runs and immense waste of training time. Exploring how to best tackle this problem could increase the optimization.

A common theme for developing tools to predict TCR specificity to the epitopes is to improve encoding of input data e.g. as observed in the tools developed using sequence similarity [14], k-mer sequence features [15], convolutional models [16], utilising amino acid physicochemical properties [17], learning a non-parametric function [7] or enhancing sequence feature into a high-dimensional space [8], utilising molecular structures [18] or pre-training BERT model on TCR data [19]. Generally, these tools train a binary classification model to exhibit epitope specificity among a background of epitopes. However, our study provided an alternative approach: a multi-class classification model towards TCR classification. Only DeepTCR included a multi-class setting, but this was not employed for the entire TCR data, only for the GAG TW10 epitope family (with ten variants). To the best of our knowledge, the multi-class approach has never been utilised in such a fashion because of scarcity of TCR data or lacking computational resources. We could overcome these limitations by utilising pre-trained transformers, trained on large corpus of protein sequence data.

We showed that the ProtXLNet transformer with the Embedding setting outperformed the previously known sequence-based and deep learning-based algorithms. However, a recently published transformer-based algorithm, i.e. TCR-BERT, was not included. TCR-BERT used two different BERT models (one for beta sequences and one for alpha sequences) and combined their embeddings for the classifier block. It does not exploit additional information, like we do on the gene usage. Despite not being able to make a head-to-head comparison with TCR-BERT, we assessed the comparisons based on the AUCROCs reported by the TCR-BERT method, but that could only be done for the CMV-pp65-NLVPMVATV class. The TCR-BERT approach resulted in an AUCROC of 0.837 (using both TRA and TRB sequences), whereas our modified ProtXLNet model had an AUCROC of 0.86 (using only TRB sequences). TCR-BERT utilized the entire 81K entries from VDJdb to pre-train a BERT model as opposed to us. We used pre-trained transformer models build from proteins in either Uniref or BFD, comprising 216 million and 2.122 billion protein sequences. For fine-tuning the classification head, we utilised only the VDJdb data with a confidence score of more than 0. TCR-BERT fine-tuned TCR specificity for binding to a positive class (epitope). Nevertheless, a more in-depth comparison of the two approaches could provide greater insight into which approach (fine-tuning or pre-training) is better suited for TCR data.

Although our approach demonstrates the potential of transformers by fine-tuning CDR3 sequences and how additional features can be juxtaposed cohesively, some limitations remain. Problematic events such as hallucination and catastrophic forgetting [20] were not evaluated. These problems are prevalent in deep neural networks and contribute significantly to misclassification. Hallucination occurs in generated sequence at the last transformer block, leading to unlikely content. Predominantly, hallucination [21,22] often occurs when generating natural languages. As we just explored classification through transformers and not generation of sequences, we don’t expect to suffer from hallucination. Catastrophic forgetting is prevalent in transformers where the weights learned during pre-training get overridden during fine-tuning. Therefore, we employed early stopping and we lowered the learning rate to prevent pre-trained knowledge from being overridden (or diminished) during fine-tuning.

Taken together we have shown the potential of transformers and multi-class learning for TCR-epitope pair prediction. The comparison with pre-existing prediction methods for TCR-epitope pairs is performed by comparing the AUROC presented in their studies which could be a limiting factor as the underlying data can have a little variation. Although transformers outperformed other methods, fine-tuning of the hyper-parameters can further enhance the performance of transformers.

## Supplementary Text

### Biology behind antigen recognition by T-Cell

Understanding T cell receptors (TCRs) relates directly to the understanding of mechanisms involved in the adaptive immune system. While adaptive immunity involves both B and T cells, we will focus on T cells which will provide us insight into cell-mediated adaptive immunity.

#### Antigen Presentation and MHC Restriction

Presenting peptides to the TCRs is done through a class of cells known as antigen-presenting cells (APC) and is presented through Major Histocompatibility complexes (MHC); this process is termed antigen presentation. Degradation of antigen is done inside APC; this degradation is undertaken through two different pathways depending on if MHCI or MHCII is expressed on the APC. The affinity of degraded peptide (or epitope) to the MHC dictates which peptide will be presented by the MHC and also to which T-Cell (CD4+ or CD8+) (Zareie, Farenc, & La Gruta, 2020).

#### Antigen Recognition

TCR expressed on either CD8+ or CD4+ T cells binds to MHC for antigen recognition by their respective T cells. T cells’ clonal nature dictates the unique binding site on its TCRs and hence its specificity. T-Cell consists of Variable (V) and Constant (C) region; it is on V region where the sequence diversity is the most concentrated and is present on both Alpha and Beta chain of TCRs. TCR genes undergo V, D and J gene rearrangement providing TCR with high diversity.

#### Gene rearrangement

Although Gene rearrangement provides TCR diversity expected at 10^18^ in humans and 10^15^ in mice, it is not random at all, and thus regulation is governed by multiple factors (Attaf, Huseby, & Sewell, 2015) (**Figure 1**). TCR beta locus comprises 46 V gene segments, followed by two groups of D, J and C gene segments (D1, J1, C1 or D2, J2, C2). D to J recombination first occurs between a D gene and one of J gene of first group or D and J gene of the second group; followed by V to the newly rearranged D and J gene. V(D)J recombination of TCR genes plays a vital role in governing the diversity of a repertoire (all unique TCR within an individual’s immune system). Additionally, random insertion and deletion of nucleotides at junctions of V, D and J give rise to hypervariable regions, known as complementarity determining regions.

**Figure 1:**
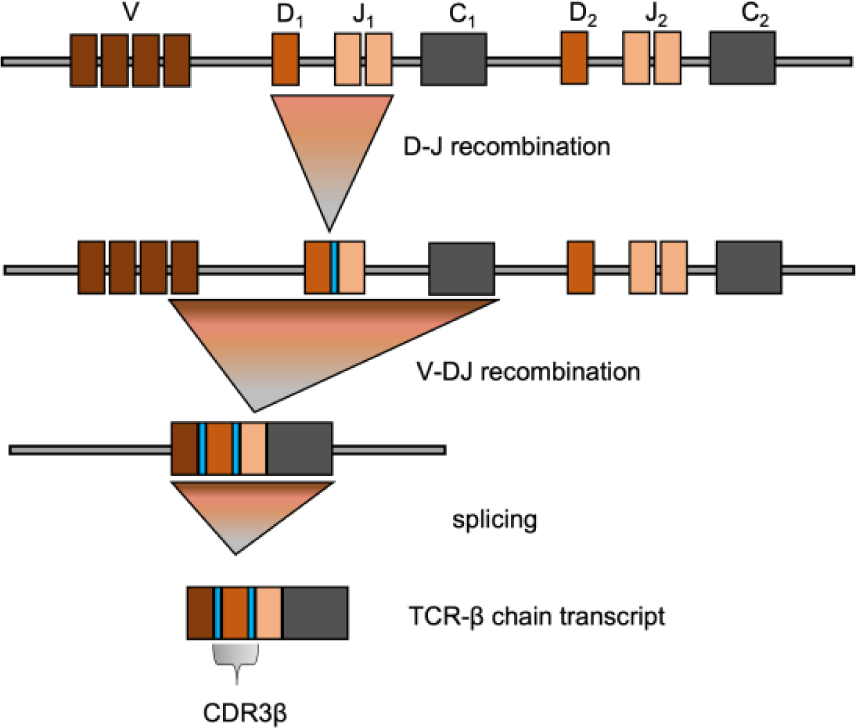
TCRβ gene rearrangement and structure. Overview of V(D)J recombination. Region in blue indicates junctional sites which are assembled by random additional and deletion. CDR3β region shown in grey on TCR protein.

#### Complementarity-determining region on TCR

Complementarity-determining regions are present at the junction of V and D segment of TCRs Fig.8. This region of TCR interacts the most with both MHC and epitope for recognition, which is due to its hypervariability. Two types of TCR, i.e., TCRAB and rarely TCRGD are mounted by the adaptive immune system for an immune response. These receptors (at protein level) are generated from two different chains, i.e., TCRAB from TCRA and TCRB and TCRGD from TCRG and TCRD. Each of these four receptors is generated from the recombination of Variable (V), diversity (D) and Joining (J) genes from their individual loci. Several V, D and J genes for TCRB and TCRD and V and J genes for TCRA and TCRG are distributed over a long stretch of chromosomes 7 and 14 in the human genome. Any of these V, D and J genes can be selected for each locus and any of the chains, i.e., TCRA/TCRB can be randomly selected to generate diverse receptors to raise an efficient immune response against infections. Overall, 10^16^ receptors can alone be generated by the V(D)J recombination events. Apart from V(D)J recombination, another factor that adds to the diversity of these receptors is the addition of nucleotides on both sides of D gene during V(D)J recombination. This region is known as complementarity determining region 3 (CDR3), which is fundamental in interacting with and recognising the antigen. Determining binding specificity to an antigen thus helps us to assess the immune system’s ability to engage with pathogens and also to evaluate the response undertaken by it.

### Transformers

Transformers are a type of Sequence to Sequence (or Seq2Seq) model that transforms a source sequence’s representation to a representation of a target sequence. Machine translation (also known as sequence transduction) which were widely performed using LSTM (long-short-term-memory) models or RNN (recurrent neural networks) models, had limitations. Limitations of fixed sequence length in traditional methods (and even the bottleneck between encoder and decoder architectures) were then overcome with the introduction of Transformers. Transformers retained the encoder-decoder architecture and introduced a new form of attention mechanism, self-attention, which outperformed standard practices.

#### Encoder-Decoder Architecture

Given a source sequence to be translated to a target sequence, the encoder would generate vector representation for each word in the source sequence, and the decoder would then read these vectors and generate words in the target sequence. Certain limitations were discovered; translating word to word doesn’t account for languages written from left to right or with different grammar compositions. Subsequently, to reduce computational resources instead of encoding the entire source sequence, a need to focus on relevant words arise.

Recurrent Neural Network (Bahdanau, Cho, & Bengio, 2014) would solve the former issue by including a weighted sum of preceding and succeeding words in a sequence whilst encoding a word. It employed RNN to encode each word, and a decoder would then decode from this representation. With the introduction of LSTMs (Sutskever, Vinyals, & Le, 2014) long-range dependencies for a word were also accounted for.

Ditching recurrent and convolutional layers was Transformers, replacing it with an attention mechanism. Transformers introduced encoding through self-attention, which would amplify the contribution of relevant words and diminish the contributions of irrelevant words while encoding a vector representation for a word. All this whilst retaining encoder-decoder architecture (there are encoder-only architectures like BERT).

#### Attention, Self-attention and multi-head attention

Origins of attention lie in the field of psychology, where the observations in behavioural patterns were attributed to where the brain was paying attention. The brain preserves computation resources by paying attention to only crucial details to reach an answer (Lindsay, 2020), which can be mimicked by utilising a weighted sum of relevant words while encoding a word. Mathematically formulating this “flexibility” in a neural network is known as the attention mechanism. Global and Local attention are some of the early attention mechanisms which were applied in computer vision (Luong, Pham, & Manning, 2015; Xu et al., 2015). With the introduction of transformers, self-attention came into the limelight.

While traditional attention mechanisms would include the contribution of surrounding words blindingly, self-attention would also include the position of the word to introduce the sense of context in encoding a word. In summary, similar words in different positions in a sequence would get different encoding.

Fig. 9 shows encoding the second word in the output sequence. Three different representation of input is used to calculate self-attention. Learning the weights for every three representations is equivalent to learning self-attention. Query and Key are used to compute scores, which undergoes some processing before taking a dot product with the Value representation. This dot product will enhance values in the Value matrix, which corresponds to higher relevance and hence augment the effect of those words into the resulting representation of the word. This can be done multiple times in parallel for a single word, termed as multi-headed attention. Multi-headed attentions allow transformers to accommodate relevance from multiple positions in a sequence.

#### Transfer Learning and Transformers architectures

Training a transformer involves the concept of transfer learning; in transfer learning, we divide training a model into two different parts: pre-training and fine-tuning. Pre-training involves two tasks mask language modelling and Next Sentence prediction. In the case of ProtTrans, only mask language modelling was utilised. The weights learned in pre-training are fine-tuned on downstream tasks, consequently saving time and resources in training a transformer from the beginning and prime advantage of transfer learning. Additionally, different Transfer models employ different approaches for pre-training tasks, which is motivated by their methodology.

There are two architectures utilised in this work, Auto-encoder and Auto-regressive. BERT, Albert, Electra are auto-encoder models with only encoders and no decoders. XLNet is an autoregressive language model. While BERT learns bidirectional language modelling, ALBERT (A lite BERT) is a more efficient version of BERT with parameter sharing among different encoder layers. Both of them use the same mask language modelling. On the other hand, ELECTRA utilises Generator-Discriminator based approach; the discriminator then detects corrupted tokens generated by the generator during masked language modelling. The discriminator trained is then used for fine-tuning on downstream tasks.

While auto-encoder models reconstruct original data during mask language modelling. They lack obvious information; masked token (it is a type of special token signified as MASK by tokeniser) will not be seen in downstream tasks, dependency learnt for the masked token (and the original token) is then not transferred (or would never be needed); termed as the pretrain-finetune discrepancy. The pretrain-finetune discrepancy is addressed in XLNet. XLNet is an auto-regressive model which implements two-stream self-attention as a means to address both forward and backward dependencies as well as to address pretrain-finetune discrepancy. Providing different factorisation orders during permutation language modelling enables the model to gather positional information from a possible position for a given token.

On the other hand, two-Stream self-attention uses additional self-attention to isolate the content while the model learns positional information (or context) during pre-training (Figure 2). Any difference between XLNet and BERT is due to this very difference in pre-training objective, which helps it retain a more significant number of dependencies than BERT. In summary, XLNet is a bidirectional transformer similar to BERT but utilises permutation language modelling.

**Figure 2:**
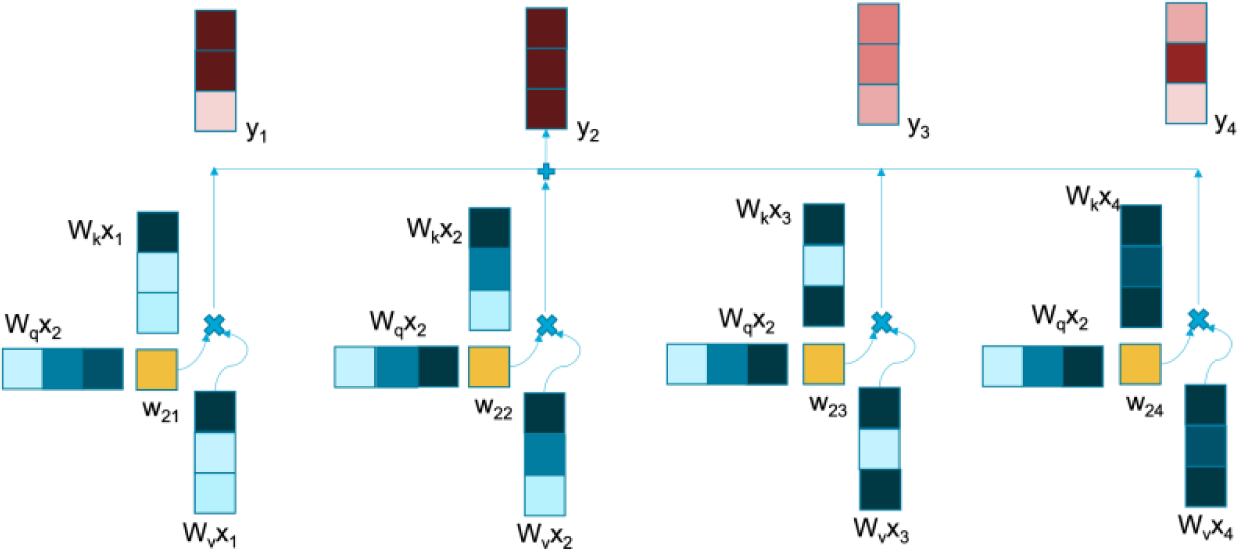
Self-Attention in Transformers. Overview of self-attention computation for encoding y2. Two representations (Query and Key) of a word are used to compute scores (shown in yellow) which enhances or reduces effect of a word (multiplying by Value). Resultant effect is summed across all words to encode a single word (y2)

---

**Supplementary Figure 1:**
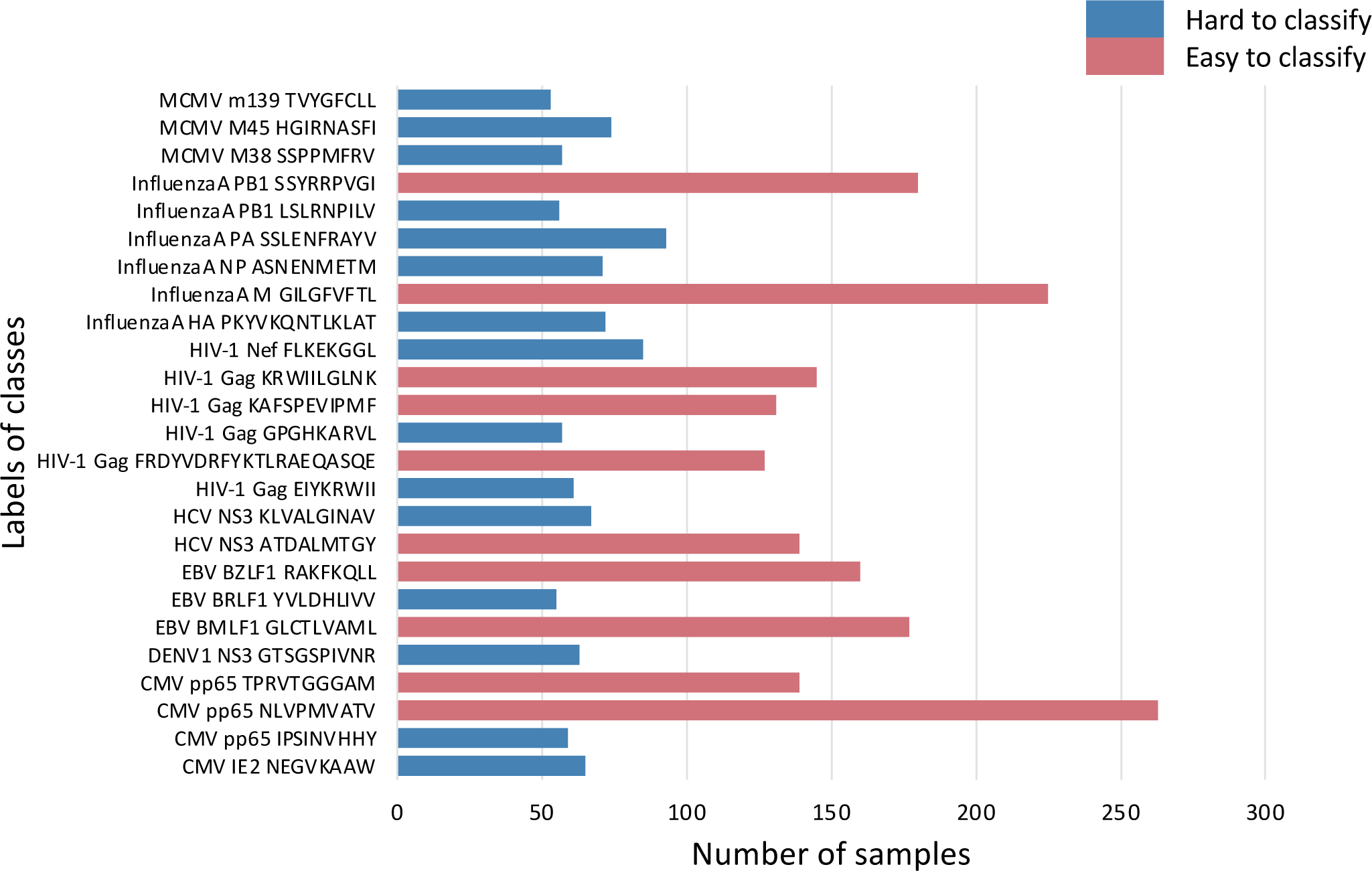
25 Epitope labels used as classes for the classification of TCR to epitopes after filtering steps as mentioned in Table 1. The classes are colored blue if the epitope is considered as hard to classify (<100 samples) and red for easy to classify labels (>100 samples).

**Supplementary Figure 2:**
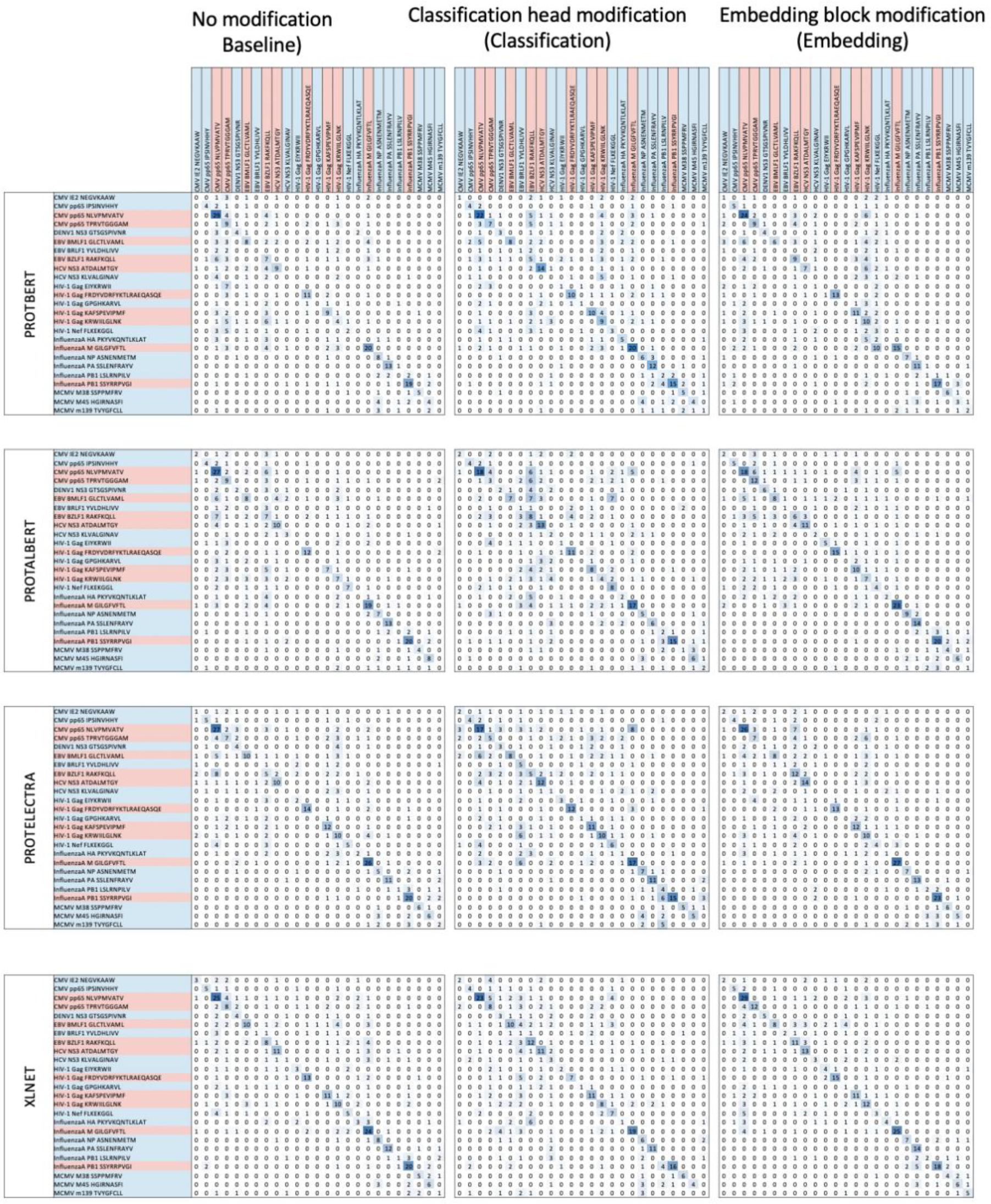
Confusion matrix of the performance of the four optimized transformers for all the three methods. The labels are colored based on their classification i.e. hard (blue) or easy (red) to classify. The matrix itself is colored from low (white) to high (blue) correctly classified labels.

**Supplementary Figure 3:**
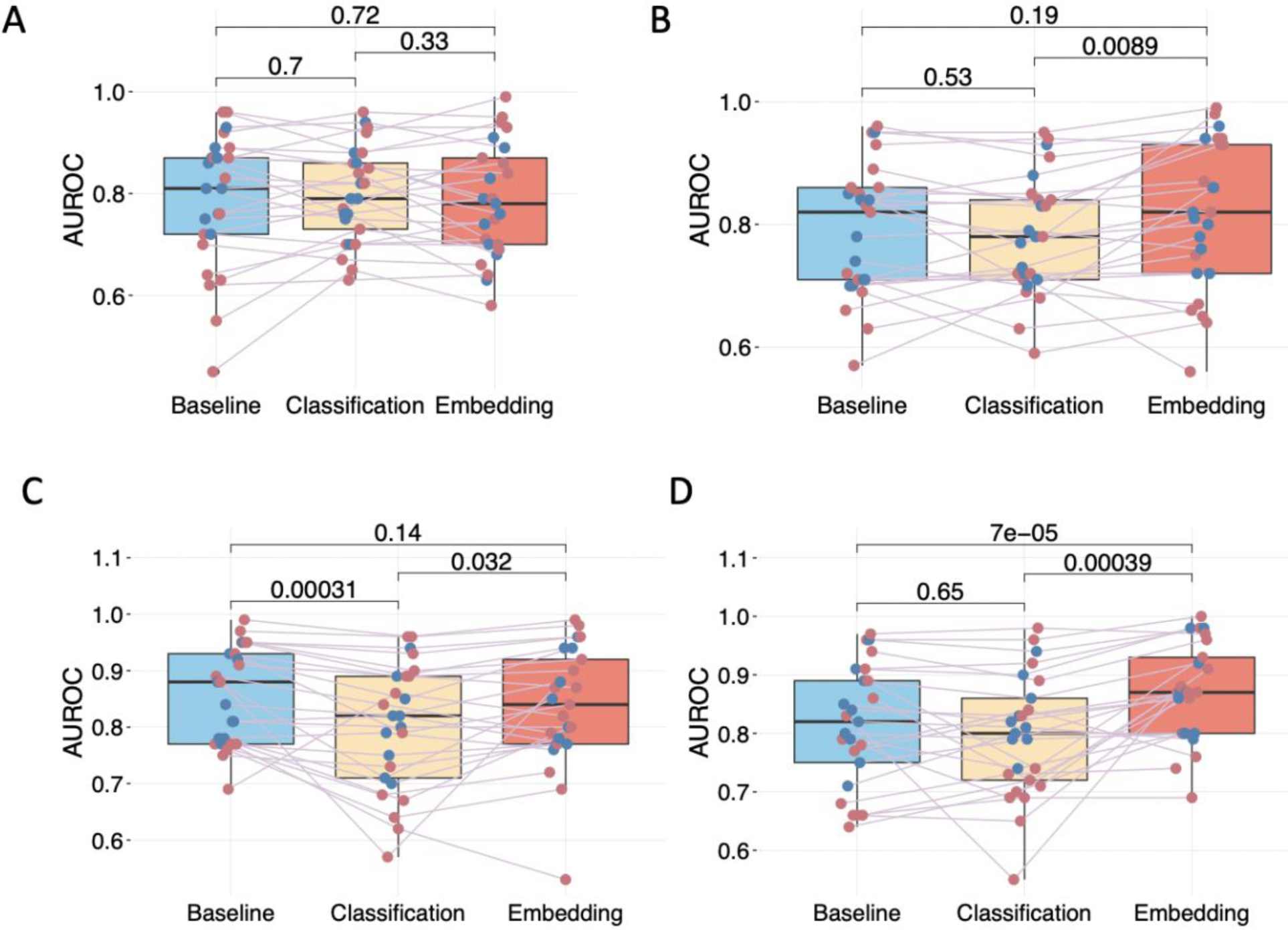
The comparison of the AUC values of the different transformer models for all the three methods A) ProtBert; B) ProtAlbert C) ProtElectra; and D) ProtXLNet. The points are colored based on the classification of the epitope classes i.e. hard (blue) or easy (red) to classify. The p values are indicated for the individual comparisons.

**Supplementary Figure 4:**
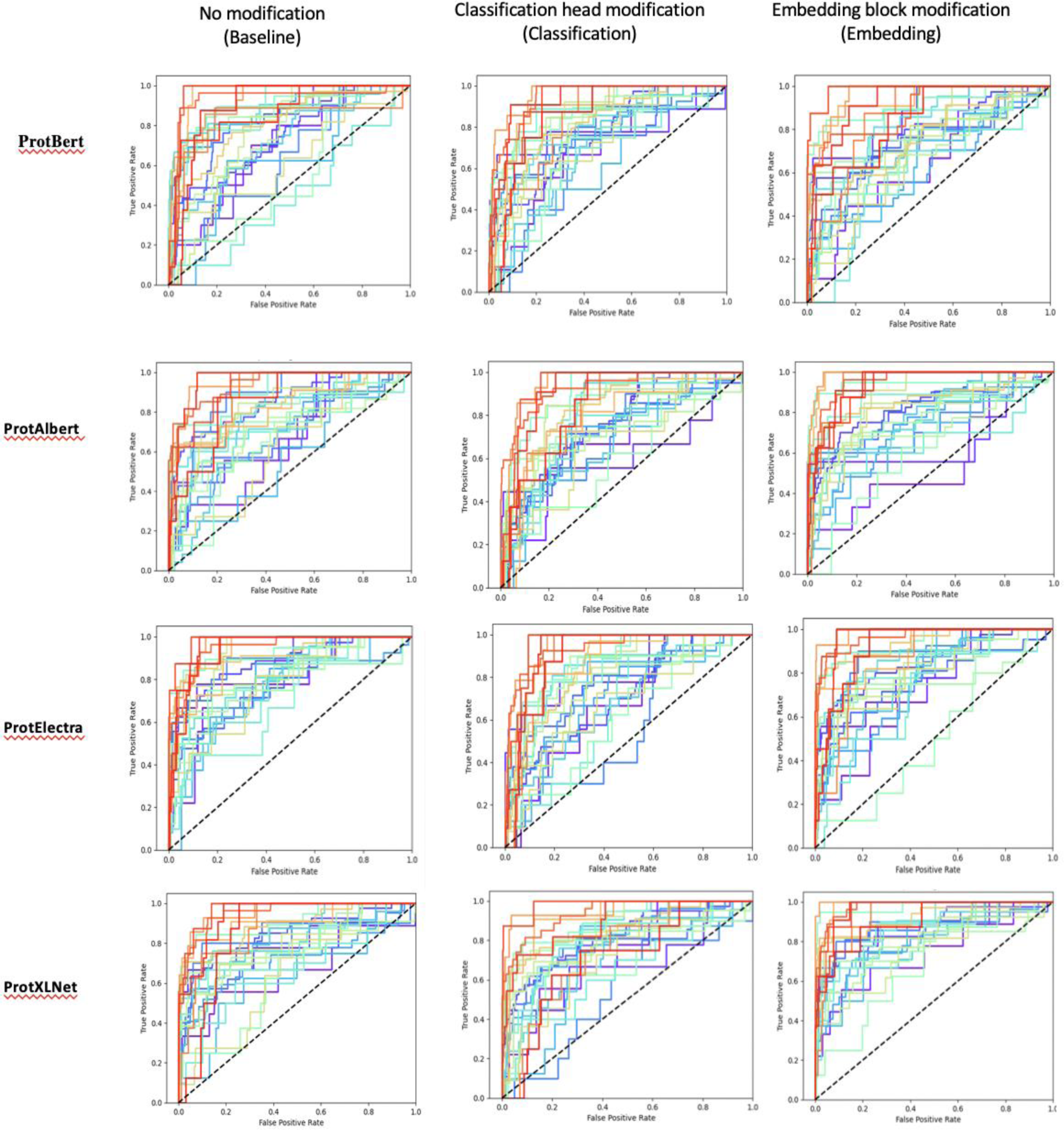
ROC plots of the performance of the 25 classes for four transformers for all the three methods. The labels are highlighted based on their classification i.e. hard (blue) or easy (red) to classify.

**Supplementary Table 1:**
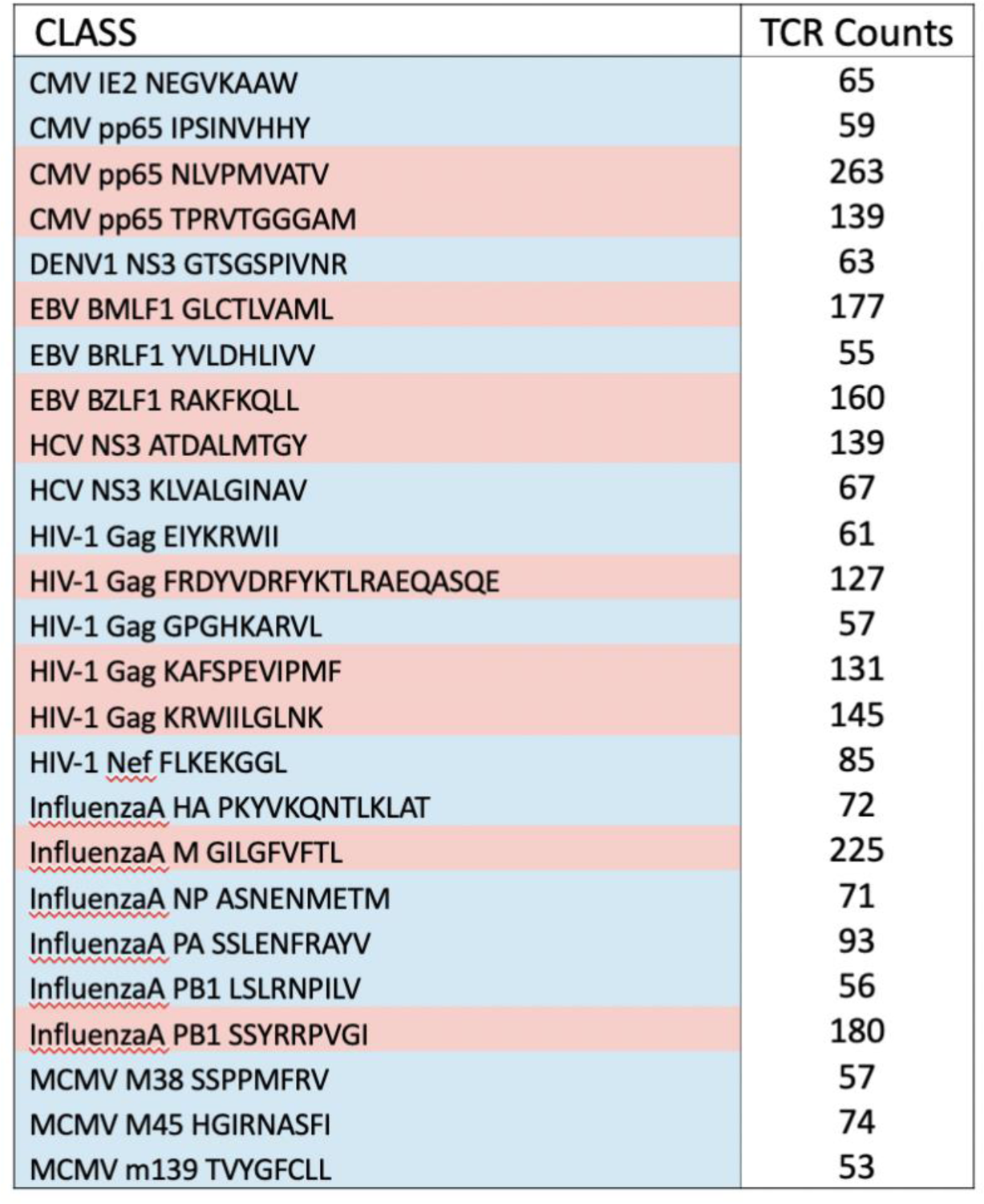
The 25 classes and corresponding count of TCRs. Each class comprise of the organism (e.g. CMV, DENV), protein name (e.g. IE2, pp65), and epitope protein sequence (e.g. NEGVKAAW, IPSINVHHY). The classes are highlighted based on their classification i.e. hard (blue) or easy (red) to classify.

**Supplementary Table 2:**
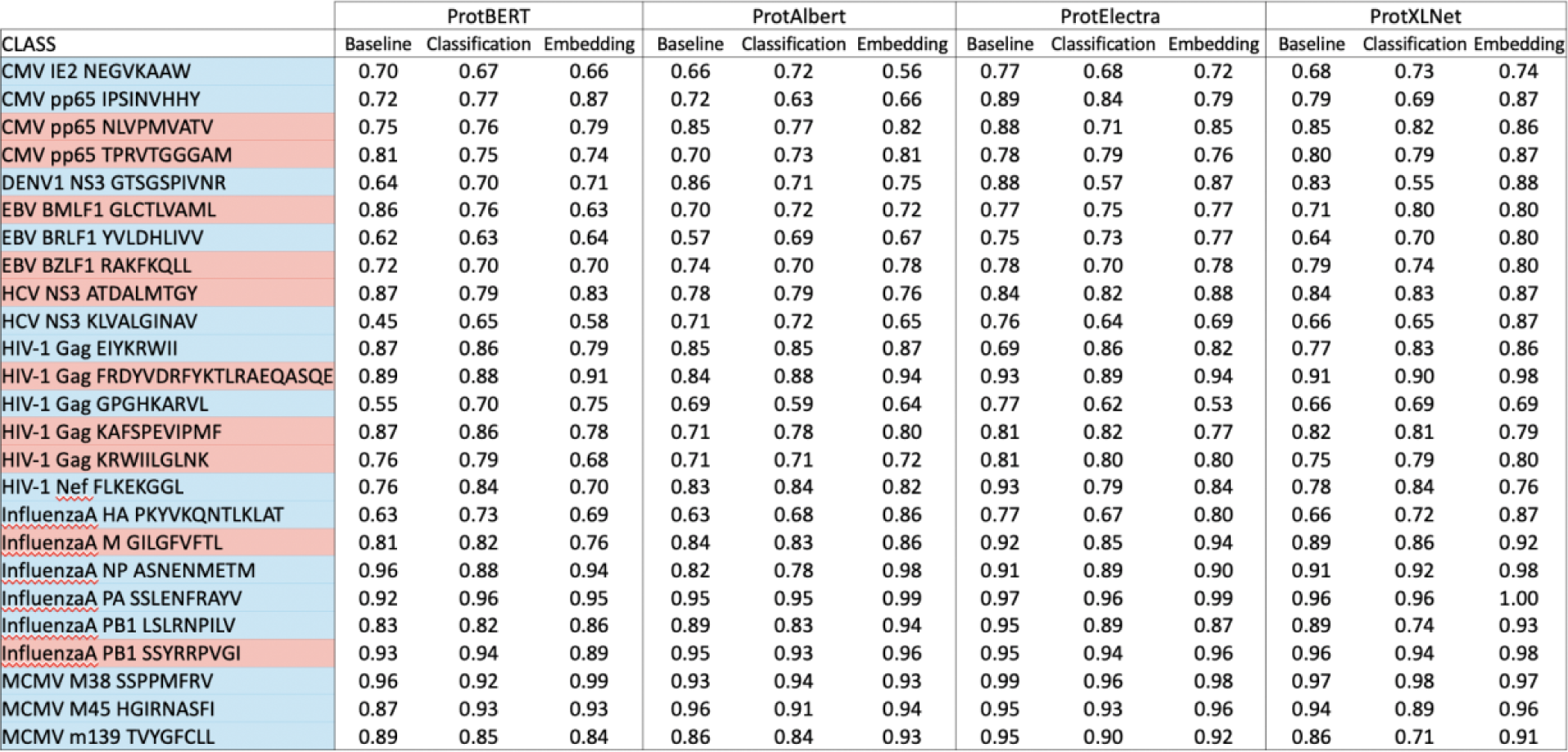
The AUC values corresponding to the ROC plots in Supplementary Figure 6. The classes are highlighted based on their classification i.e. hard (blue) or easy (red) to classify.

**Supplementary Table 3:** The AUC values of the publicly available tools compared in the study.

| CLASS | TCRGP | TCRDist | DeepTCR |
| --- | --- | --- | --- |
| CMV IE2 NEGVKAAW | NA | NA | NA |
| CMV pp65 IPSINVHHY | 0.85 | 0.80 | 0.78 |
| CMV pp65 NLVPMVATV | 0.89 | 0.79 | 0.87 |
| CMV pp65 TPRVTGGGAM | 0.91 | 0.83 | 0.86 |
| DENV1 NS3 GTSGSPIVNR | 0.86 | 0.88 | 0.88 |
| EBV BMLF1 GLCTLVAML | 0.93 | 0.86 | 0.91 |
| EBV BRLF1 YVLDHLIVV | 0.68 | 0.73 | 0.56 |
| EBV BZLF1 RAKFKQLL | 0.89 | 0.76 | 0.78 |
| HCV NS3 ATDALMTGY | 0.82 | 0.75 | 0.85 |
| HCV NS3 KLVALGINAV | 0.68 | 0.70 | 0.70 |
| HIV-1 Gag EIYKRWII | 0.77 | 0.54 | 0.77 |
| HIV-1 Gag FRDYVDRFYKTLRAEQASQE | 0.75 | 0.77 | 0.66 |
| HIV-1 Gag GPGHKARVL | 0.70 | 0.69 | 0.70 |
| HIV-1 Gag KAFSPEVIPMF | 0.89 | 0.84 | 0.86 |
| HIV-1 Gag KRWIILGLNK | 0.84 | 0.70 | 0.66 |
| HIV-1 Nef FLKEKGGL | 0.80 | 0.67 | 0.83 |
| InfluenzaA HA PKYVKQNTLKLAT | 0.71 | 0.75 | 0.60 |
| InfluenzaA M GILGFVFTL | 0.88 | 0.83 | 0.89 |
| InfluenzaA NP ASNENMETM | 0.90 | 0.84 | NA |
| InfluenzaA PA SSLENFRAYV | 0.94 | 0.89 | NA |
| InfluenzaA PB1 LSLRNPILV | 0.82 | 0.80 | NA |
| InfluenzaA PB1 SSYRRPVGI | 0.91 | 0.90 | NA |
| MCMV M38 SSPPMFRV | 0.92 | 0.92 | 0.84 |
| MCMV M45 HGIRNASFI | 0.89 | 0.85 | 0.90 |
| MCMV m139 TVYGFCLL | 0.72 | 0.66 | 0.79 |

